# Large-scale organization of the hand action observation network in individuals born without hands

**DOI:** 10.1101/305888

**Authors:** Gilles Vannuscorps, Moritz Wurm, Ella Striem-Amit, Alfonso Caramazza

**Affiliations:** Department of Psychology, Harvard University, Cambridge, MA, 02138, USA Center for; Mind/Brain Sciences, Università degli Studi di Trento, Mattarello, 38122, Italy; Faculty of Psychology and Educational Sciences and Psychological Sciences Research Institute, Université catholique de Louvain, Louvain-la-Neuve, 1348, Belgium

## Abstract

The human high-level visual cortex comprises regions specialized for the processing of distinct types of stimuli, such as objects, animals, and human actions. How does this specialization emerge? Here, we investigated the role of sensorimotor experience in shaping the organization of the action observation network as a window on this question. Observed body movements are frequently coupled with corresponding motor codes, e.g. during monitoring one’s own movements and imitation, resulting in bidirectionally connected circuits between areas involved in body movements observation (e.g., of the hand) and the motor codes involved in their execution. If the organization of the action observation network is shaped by this sensorimotor coupling, then, it should not form for body movements that do not belong to individuals’ motor repertoire. To test this prediction, we used fMRI to investigate the spatial arrangement and functional properties of the hand and foot action observation circuits in individuals born without upper limbs. Multivoxel pattern decoding, pattern similarity, and univariate analyses revealed an intact hand action observation network in the individuals born without upper limbs. This suggests that the organization of the action observation network does not require effector-specific visuomotor coupling.

## Introduction

The high-level visual cortices contain a reproducible and consistent spatial arrangement of areas specialized for the processing of distinct types of stimuli such as manipulable objects (Lewis 2006), animals (Konkle and Caramazza 2013), faces (Kanwisher et al. 1997), other body parts (Downing et al. 2001), and human actions (Beauchamp et al. 2002; Caspers et al. 2010). The origin of this organization remains unclear, however.

Research on this issue in recent years affords two conclusions. First, this large-scale organization does not require visual experience: congenitally blind individuals show the stereotypical large-scale organization of domain preference (Mahon et al. 2009; He et al. 2013; Ricciardi et al. 2013; Peelen et al. 2014; Striem-Amit and Amedi 2014; van den Hurk et al. 2017). Second, there is mounting evidence that this organization draws on the large-scale connectivity pattern of these different areas to various motor-, affective– or cognitive-related downstream computational systems and, thus, reflect how the information processed by the different preference areas is to be used by the rest of the brain in the service of behavior (Mahon and Caramazza 2011; Saygin et al. 2011; Bracci et al. 2012; Simmons and Martin 2012; Kravitz et al. 2013; Hutchison et al. 2014; Hannagan et al. 2015; Heimler et al. 2015; Bi et al. 2016; Saygin et al. 2016; Konkle and Caramazza 2017; Stevens et al. 2017).

What remains unknown, however, is whether this organization emerges from repeated functional coupling between the area of specialization within the high-level visual cortex and their downstream computational systems (Johnson 2011; Hannagan et al. 2015), possibly through Hebbian learning mechanisms (Hebb 1949). Alternatively, its emergence may be guided by a genetically determined self-organizing process independent from any type of experience (Mahon and Caramazza 2011; Hannagan et al. 2015; Saygin et al. 2016). To date, only a few studies have provided data relevant to this question, and their results diverge. In favor of a critical role for experience, it has been shown that the emergence of the brain’s typical area of specialization for letters requires reading experience (Baker et al. 2007; Dehaene et al. 2010; Saygin et al. 2016). By contrast, the neural representation of tools and hands appears to be virtually the same in congenitally blind individuals (Peelen et al. 2013; Kitada et al. 2014; Striem-Amit and Amedi 2014), individuals born without hands (Striem-Amit et al. 2017), and in the typically developed population (Bracci et al. 2012). Here, we investigated the role of sensorimotor experience in shaping the organization of the action observation network as a window on this question.

A bilateral network of three main brain regions commonly referred to as the “action observation network” (AON) – the lateral occipitotemporal cortex (LOTC), the inferior parietal lobule (IPL) and the ventral premotor cortex (PMv) – has been consistently reported to preferentially respond to observed human actions than other animals’ or objects’ shape and movements (Grezes et al. 2001; Grossman and Blake 2002). This network forms a bidirectionally connected functional circuit translating observed actions into the corresponding motor codes, and vice versa. Viewing an action executed with a specific set of muscles, for instance, activates a representation of the corresponding muscles in the observer’s brain (Maeda et al. 2002; Fadiga et al. 2005). In the opposite direction, the execution of unseen hand and arm movements has been shown to engage the LOTC and the IPL (Astafiev et al. 2004). Thus, this network’s specialization for human actions could emerge through repeated functional coupling between the three regions of the AON (Wilson and Knoblich 2005; Heyes 2010; Casile et al. 2011) in the service of essential functions such as imitation (Buccino et al. 2004), observational learning (Mattar and Gribble 2005), and the visual control of one’s own body movements (Oztop and Arbib 2002; Wolpert et al. 2003). To test this possibility, we explored how people born without hands (individuals with dysplasia; IDs) represent hand and foot actions. We tested if hand action observation can be decoded in the IDs in the same locations as in controls. If the typical large-scale organization and functional properties of the network for action representation emerges through repeated effector-specific functional coupling between the three regions of the AON, then, hand action representation should not be found in the AON in the IDs. If, however, the large-scale functional organization of the AON does not require effector-specific visuomotor coupling, then, the IDs should represent hand actions in the AON.

## Materials and Methods

### Participants

Five individuals (two males) born with severely shortened or completely absent upper limbs (participants with bilateral upper limb dysplasia, IDs, see Figure 1) of ototoxic (in-utero thalidomide exposure, ID2), genetic (ID5) or unknown origin and nine typically developed control participants (TDs) matched for age (no group difference; p < 0.29) participated in the experiment. No participant had a history of psychiatric or neurological disorder. All the IDs had developed fine motor skills of the feet and used their feet for many typical hand-related actions of daily life (e.g., opening or closing doors; for more information about IDs’ foot use see Vannuscorps et al. 2014; Striem-Amit et al. 2017) and none had history of phantom limb sensations or movements (tested as in Vannuscorps and Caramazza 2016). Two TDs were excluded from the sample due to poor behavioral performance (error rate exceeding the group mean by > 2 SDs) in the experimental task. All participants gave written informed consent prior to the study, which was approved by the Committee on the Use of Human Subjects, Harvard University.

**Figure 1.**
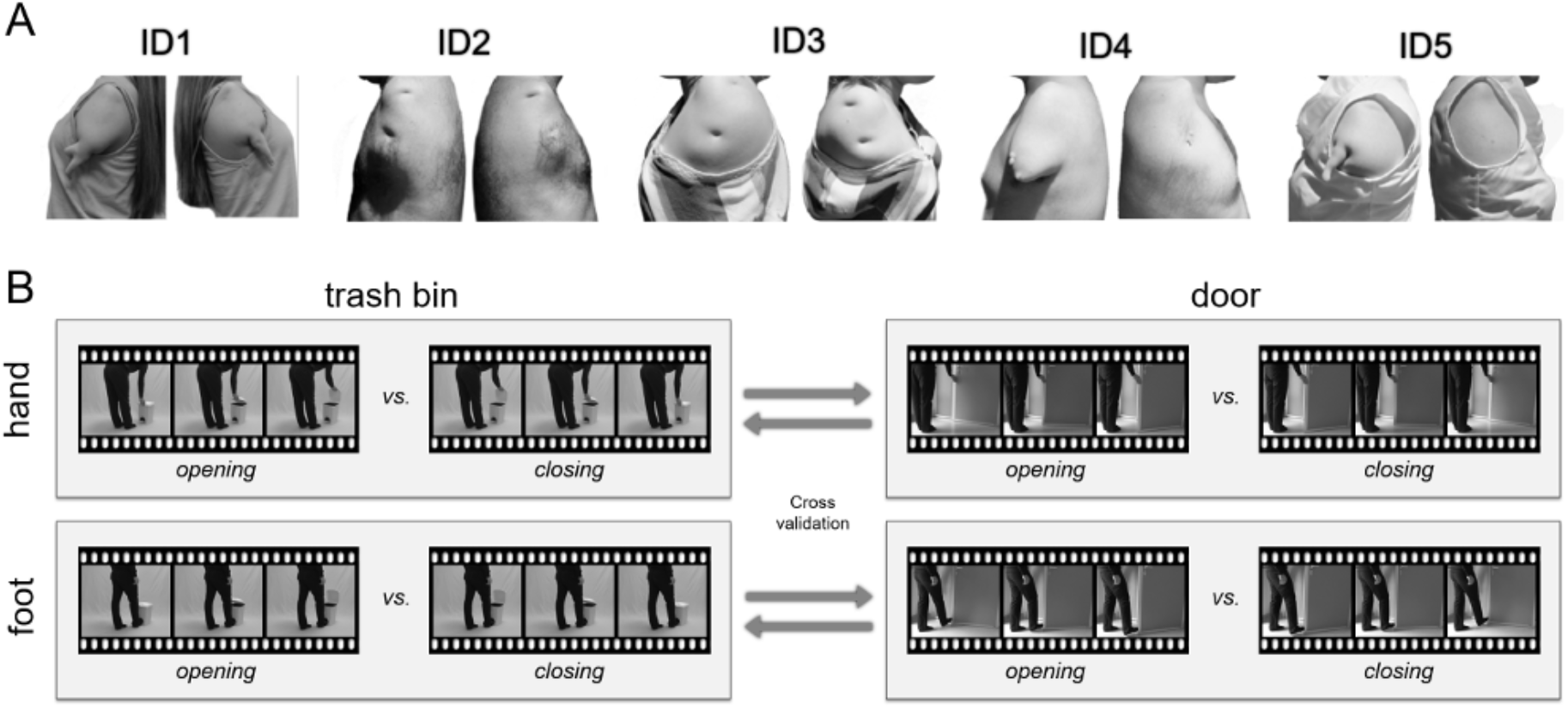
(A) Upper limb extremities of the individuals with upper limb dysplasia (IDs) included in the study. (B) Action decoding scheme and examples of the experimental conditions. In the across object decoding, a classifier was trained to discriminate between opening vs. closing trash bins and tested on opening vs. closing doors, and vice versa (for each effector separately). The decoding thus targeted action representations that generalize across the objects involved in the actions and the specific effector movements required for the actions.

### Experimental Design

The stimuli were 64 video-clips of interest and 48 video clips used as “catch trials”. The 64 video-clips of interest were 32 exemplars of “opening” and 32 exemplars of “closing” actions performed with the right hand or foot with four different exemplars of two different object types (doors and trash bins) by two different actors (see Figure 1). Representative exemplars of the video-clips can be visualized on http://www.testable.org/experiment/7/199225/start. Briefly, the actions looked as follows: the actor stood next to a door partly (about 10 cm) or largely (about 90 cm) opened with his/her hand or his/her foot positioned on the edge of the door or stood next to an opened or closed trash bin with his/her hand positioned on the edge of the trash bin’s lid or his/her foot positioned on the trash bin’s pedal and then either opened or closed the door or the trash bin entirely. These stimuli satisfied three requirements. First, they corresponded to actions regularly executed by typically developed participants with the foot or the hand. Second, the two objects required very different kinematics (e.g., pushing/pulling vertically vs. horizontally). We thereby guaranteed that across object MVP decoding (training on one object and testing on the other) relied on abstract action representations that generalize across perceptual information (Wurm and Lingnau 2015; Wurm et al. 2016). Perceptual variance was further ensured by the use of different actors and exemplars of the two objects shown from a left or right viewpoint. Third, the upper limb actions corresponded to a movement impossible to execute with the lower limbs. The feet, for instance, allow neither grasping a bin lid with a precision grip opposing the fingertips nor grasping a door panel with a palm power grip. Thus, these actions were possible to execute by the IDs, but with different movement parameters. The 48 video clips used as “catch trials” were similar video-clips in which the action was incomplete, i.e., the door or trash bin was touched by the hand or foot but kept open or closed, respectively. During the experiment, these stimuli were back-projected onto a screen via a liquid crystal projector and viewed through a mirror mounted on the head coil. All videos were in grayscale, had a duration of 2 s and had a resolution of 768 × 576 pixels.

Each participant was scanned in a single session consisting in one anatomical scan followed by 4 functional scans (runs) during which they viewed pictures of tools, non-manipulable objects, hands and feet (these data are presented in Striem-Amit, Vannuscorps & Caramazza, 2017) and then 6 other functional scans (runs) constituting the experiment reported herein. Each run started with a 10-s fixation period, ended with a 16-s fixation period, and was constituted of 80 events lasting 2 seconds and followed by a 1s fixation period. These 80 events always comprised the 64 video-clips of interest (2 actions × 2 actors × 2 effectors × 2 objects × 4 exemplars), 8 catch trials and 8 null events, mixed in a different, first-order counterbalanced (Aguirre 2007), order in each run. Hence, there was a total of 8 trials for each combination of action, effector and object in each run, for a total of 48 trials in the whole experiment. Participants were instructed to attentively watch the movies and respond (by foot response) to the catch trials in which the action was incomplete, i.e., the door or trash bin was touched by the hand or foot but kept open or closed, respectively. The data from these trials were excluded from further analysis. Stimulus presentation, response collection, and synchronization with the scanner were controlled with Presentation^®^ software (Neurobehavioral Systems, Inc., Berkeley, CA, www.neurobs.com).

### Data acquisition

Functional and structural data were collected using a Siemens Tim Trio 3-T scanner at the Center for Brain Science at Harvard University and a 16-channel birdcage head coil. Functional images were acquired with a T2*-weighted gradient echo-planar imaging (GE-EPI) sequence that employed multiband RF pulses and Simultaneous Multi-Slice (SMS) acquisition (Moeller et al. 2010; Setsompop et al. 2012) with fat suppression. The SMS-EPI acquisitions used a modified version of the Siemens WIP 770A. Acquisition parameters were a repetition time of 2 s, an echo time of 28 ms, a flip angle of 80°, a field of view of 216 mm, and a matrix size of 108 × 108. We used 69 slices, acquired in ascending interleaved order, with a thickness of 2 mm and no gap. Slices were tilted to run parallel to the superior temporal sulcus. In each functional run, 180 images were acquired.

Structural T1-weigthed images were acquired with an MPRAGE sequence (176 sagittal slices, TR = 2530 s, inversion time = 1200 ms, FA = 7°, 256 × 256 mm FOV, 1 × 1 × 1 mm resolution).

### Preprocessing

Data were analyzed using SPM8, BrainVoyager QX 2.8 (BrainInnovation) in combination with the NeuroElf Toolbox, and custom software written in Matlab (MathWorks). The first 4 volumes were removed to avoid T1 saturation. The first volume of the first run was aligned to the high-resolution anatomy (6 parameters). Data were 3D motion corrected (trilinear interpolation, with the first volume of the first run of each participant as reference), followed by slice time correction, high-pass filtering (cutoff frequency of 3 cycles per run), and isovoxel transformation to 3 x 3 x 3 mm resolution. Spatial smoothing was applied with a Gaussian kernel of 8 mm FWHM for univariate analysis and 3 mm FWHM for MVPA (Wurm and Lingnau 2015; Gardumi et al. 2016). Anatomical and functional data were transformed into MNI space using trilinear interpolation.

### Statistical Analysis

#### Multivoxel pattern analysis (MVPA)

MVPA was carried out using linear discriminant analysis (LDA) classification as implemented in the CoSMoMVPA toolbox (Oosterhof et al. 2016). Design matrices contained 16 predictors reflecting the action conditions (8 actions x 2 exemplars), 2 catch trials predictors, and 6 predictors resulting from 3D motion correction. Each predictor was convolved with a dual-gamma hemodynamic impulse response function (Friston et al. 1998). Each trial was modeled as an epoch lasting from video onset to offset (2 s). The resulting reference time courses were used to fit the signal time courses of each voxel. Beta weights of experimental conditions were estimated on the basis of 4 trials per condition and run resulting in two beta estimates per action condition and run. The 4 trials were selected from either the first half or the second half of each run. Because the 4 trials showed different instantiations of the same action (different viewpoints, actors, and object exemplars), the MVPA targeted action representations that generalize across these factors (Wurm et al. 2017). In total, this procedure resulted in 12 beta maps per action condition (6 runs x 2 exemplars). Searchlight-based MVPA (Kriegeskorte et al. 2006) was performed in volume space using spherical ROIs with a radius of 12 mm (Wurm and Lingnau 2015).

The following steps were done for each participant and searchlight ROI separately. Within each individual ROI (221 voxels on average), beta weights were extracted resulting in 12 beta patterns per condition. Beta patterns were then entered into MVPA. The “across object” decoding targeted action representations that generalize across object categories and thus are independent of the visual features of the stimuli. Using a leave-one-run-out cross validation scheme, a classifier was trained to discriminate between the beta patterns of opening and closing a door with the one effector (i.e., the hand or the foot) and tested on the beta patterns of opening and closing a trash bin with the same effector. The same classification procedure was done vice versa, i.e., the classifier was trained on opening and closing a trash bin and tested on opening and closing a door, and the decoding accuracies were averaged across the two generalization directions and across the 6 iterations of the cross-validation. MVPA thus resulted in one mean decoding accuracy value per effector, ROI, and subject. The “across effector” decoding targeted action representations that generalize across the effector that is used for the action. In this decoding scheme, a classifier was trained to discriminate between opening vs. closing trash bins with the hand and tested on opening vs. closing trash bins with the foot, and vice versa (Figure S1A). The same was done for actions involving doors; resulting accuracies were averaged across object types. The “across object and effector” decoding, targeted action representations that generalize across both the object involved in the action and across the effector that is used for the action. To this end a classifier was trained to discriminate between opening vs. closing trash bins with the hand and tested on opening vs. closing doors with the foot, and vice versa (Figure S1A). The same was done for the other possible combination (train on opening vs. closing trash bins with the foot and tested on opening vs. closing doors with the hand, opening vs. closing trash bins with the hand), and resulting accuracies were averaged. In all decoding analyses mean accuracy values were assigned to the center voxel of each searchlight sphere.

Additionally, we computed 100 chance accuracy (null) maps for each decoding analysis. Each null map was computed in the same manner as reported above, except that the target labels were randomized before decoding to generate classification measure outcomes under the null hypothesis. Labels were randomized under the constraint that the number of samples in training and testing datasets was balanced between the two classes to avoid biases due to uneven class distributions in the training and testing datasets (Nichols and Holmes 2002; Oosterhof et al. 2016). Null maps were entered into permutation tests at the group level (see below) by randomly selecting one null map per subject for subsequent group analysis (Nichols and Holmes 2002; Stelzer et al. 2013).

#### ROI MVPA

AON ROIs (12 mm radius) were defined based on coordinates from the meta-analysis for action observation of Caspers et al. (2010) [MNI coordinates; PMv: -50/9/30 (left), 52/12/36 (right); IPL: -60/-24/36 (left), 44/-34/44 (right); and LOTC: -46/-72/2 (left), 52/-64/0 (right)]. This subject-independent ROI definition was chosen to avoid group-specific ROI selection biases. Decoding accuracies were extracted from searchlight maps. For stability and normality, we averaged the decoding accuracies across voxels of a ROI (Fairhall and Caramazza 2013). For each test and ROI, we first tested whether the mean decoding accuracies of the TDs and the IDs was significantly higher than the classification expected by chance (50%). To this end we estimated the null distribution of mean decoding accuracy for each subject group, test, and ROI from resampled null maps. Resampling of each subject group was done by randomly selecting one of the 100 null maps per subject to obtain a unique combination of null maps (10000 iterations). For each iteration, ROI, test, and subject group, decoding accuracies were extracted from the resampled maps, averaged across voxels, and the mean decoding accuracy of the group was computed. We then estimated the probability that the observed true mean accuracy can occur by chance by counting the number of mean accuracies of the null distribution that are equal or greater than the true mean accuracy and divided that number by the number of all mean accuracies of the null distribution (10000). The resulting p values of each subject group were FDR-corrected (Benjamini and Yekutieli 2001) by the number of ROIs and tests (24 tests in total). Additional analyses that tested for deviant hand action decoding in the IDs were carried out only in those ROIs that showed significant above chance accuracies in the TDs or in the IDs (see Supplemental Material).

#### Whole-brain MVPA

For whole-brain analysis, individual accuracy maps of the hand and foot action decoding were entered into a one-sample t-test for each subject group to identify voxels yielding classification significantly above chance. Statistical maps were corrected for multiple comparisons using a cluster-based Monte Carlo simulation algorithm as implemented in the CoSMoMVPA Toolbox (Oosterhof et al. 2016). We used a threshold of p = 0.05 at the cluster level, and initial voxelwise threshold of p = 0.001, and 10000 iterations of Monte Carlo simulations. For visualization, maps were projected on a cortex surface of a Colin27 MRI volume as provided by the Neuroelf toolbox using BrainVoyager QX 2.8 (BrainInnovation). In a second-level analysis, we computed the interaction GROUP (TDs, IDs) x EFFECTOR (hand, foot) as reported above (ROI MVPA). This allowed us to identify voxels that are in favor of the specific hypothesis that the IDs show weaker decoding for hand vs. foot actions (indicated by positive t values; note however that positive t values may also indicate weaker decoding of foot vs. hand actions in the TDs). The maps were thresholded using a bootstrapping procedure (Nichols and Holmes 2002) using a cut-off at p = 0.05 to isolate voxels in which the observed t value based on the true assignment into TDs and IDs is higher than 95% of t values based on random assignments into TDs and IDs. To this end we computed 10000 unique random group assignments permutations not identical to the true group assignment by shuffling the group labels (TDs, IDs). For each random assignment we computed a new t map. Then we computed, for each voxel, the upper bound of the 95% confidence interval (one-sided, z = 1.6449) of the bootstrapping distribution and tested whether the t value of the true group assignment exceeds this cut-off. The resulting map was further thresholded at t(10) = 3.15 (p = 0.01) and a cluster size of 3 voxels.

#### Whole-brain decoding map similarity

To provide positive evidence that IDs represent hand actions like the TDs, we employed a multivariate similarity analysis. We reasoned that if the IDs represent hand actions in similar brain regions, and thus voxels, as the TDs, then the decoding maps of the IDs and the TDs should significantly correlate with each other. To this end, we compared the correlation of decoding maps within the TD with the correlation of decoding maps between TDs and IDs.

We first computed the mean accuracy map correlation within the TDs. To this end, the TDs were split into two groups of 3 and 4 subjects, respectively, using all permutations of possible subject combinations. For each permutation and subject group, mean accuracy maps were computed. Maps were then vectorized and correlated with each other using Pearson correlation. Resulting correlation coefficients were averaged across permutations.

To test whether there was a significant correlation within the TDs, we tested the within-TD correlation against the null distribution of correlation coefficients based on random permutation maps. Random permutation maps were generated for each participant by permuting the decoding targets (opening vs. closing) to generate decoding accuracies under the null hypothesis. For each participant, a set of 100 randomized maps was generated. Using a bootstrapping approach as described in Stelzer et al. (2013), individual maps were randomly selected from the set and combined to 10000 different group pools. For each pool, a within-TD mean correlation coefficient was computed as described above. The true correlation coefficient was then tested against the 95% confidence interval (CI) of the null distribution.

In a similar manner, between-group mean accuracy map correlations were computed. To this end, permutations of all possible subsets of 3 and 4 TDs, respectively, were computed. Likewise, permutations of all possible subsets of 3 and 4 IDs, respectively, were computed. For each subset, maps were averaged and vectorized. TD and ID subsets were then correlated with each other, so that subsets were always based on 3 TD and 4 ID maps or 4 TD and 3 ID maps. Resulting correlation coefficients were averaged across permutations. As described above, a null distribution of between-group correlations was computed, and the mean between-group correlation coefficient was tested against the 95% confidence interval (CI) of the null distribution.

#### Univariate fMRI analysis

For each participant, a general linear model (GLM) was computed using design matrices containing predictors of the 8 action conditions, catch trials, and of the 6 parameters resulting from 3D motion correction (x, y, z translation and rotation). For each participant, the contrasts hand actions vs. baseline and foot actions vs. baseline were computed. For ROI analysis, beta estimates were extracted from spherical ROIs with 12 mm radius for each contrast map and participant. AON ROIs were defined as in the MVPA reported above. Beta values were averaged across ROI voxels and entered into one-tailed one sample tests. ROIs that showed significant activations in at least the TDs or IDs alone (as indicated by FDR corrected significant beta coefficients) were further analyzed in subsequent second level analyses (see Supplemental Material).

## Results

We recorded functional magnetic resonance imaging (fMRI) data from typically developed participants (TDs) and five individuals born with no elbow, wrist, or hands (IDs, see Figure 1A) when watching video-clips of typical actions executed with the lower and upper limbs, with different objects, actors and viewing angles (Figure 1B). All participants accomplished the task with high accuracy. IDs made less errors (0.8 ± 0.3%) than the TDs (1.8 ± 0.4%), but the difference was not significant.

### Decoding of hand and foot actions in the AON

To test whether the IDs show the typical large-scale organization and functional properties of the brain areas specialized for observed human actions, we first conducted multivoxel pattern analyses of the fMRI data in the three brain regions of the AON (12 mm spherical regions of interest; ROIs) defined based on a meta-analysis on action observation (Caspers et al. 2010). For each ROI, participant, and effector shown in the video (hand and foot) we trained a linear discriminant analysis (LDA) classifier to discriminate between two actions (opening and closing) performed with one object (e.g., a door) and tested the ability of the classifier to discriminate the two actions performed with another object (e.g., a trash bin, see Figure 1B). Critically, this conservative generalization scheme made sure that the MVPA relied on higher-level representations of the actions instead of the mere visual features of the stimuli. We tested hand action decoding in the IDs using permutation testing, i.e., through estimating the probability of observed mean decoding accuracies relative to the null distribution of mean decoding accuracies generated from resampled random decoding data (see Methods for details).

The results of these analyses, shown in Figure 2, indicated significant above chance decoding accuracies for hand actions not only in the TDs’ bilateral LOTC and left IPL ROIs, but also in the IDs’ bilateral IPL and right LOTC ROIs (FDR corrected one-sided permutation tests, all *ps* < 0.008). This indicated that the large-scale functional organization of action representation in the areas of the AON does not require effector-specific visuomotor coupling. To further explore the data, we searched for possible differences between the hand action decoding in the IDs and in the TDs, and between the hand and foot action decoding in the IDs using traditional and Bayesian mixed ANOVAs and two-sample t tests. The results of these comparisons (see supplemental results and tables S1 and S2) failed to reveal any significant differences and, if anything, suggested that the decoding of hand and foot action were more likely similar than different between the groups.

**Figure 2.**
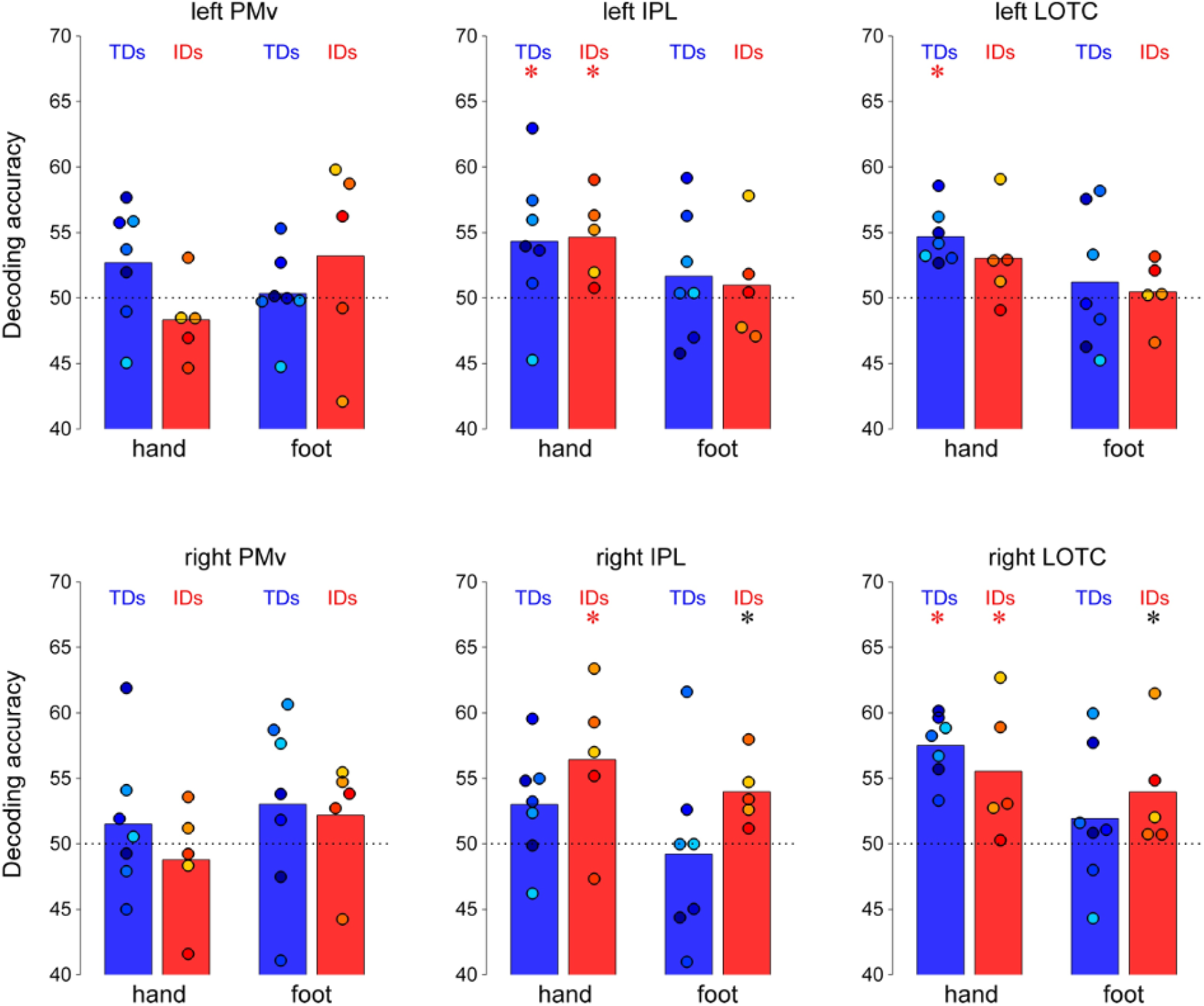
Hand and foot action decoding accuracy in the action observation network (AON). Blue and red bars represent mean decoding accuracies (50% = chance) of TDs and IDs, respectively. Superimposed colored dots indicate individual decoding accuracies. Asterisks above bars indicate significance (red: FDR corrected for the number of tests = 24; black: < 0.05 uncorrected). Both IDs and TDs showed significant hand action decoding in LOTC and IPL regions.

### Whole-brain decoding of hand and foot actions

The findings of the ROI analysis were corroborated by those of a whole brain action decoding analysis. As shown in Figure 3A, hand action decoding was strongest in LOTC and IPL for both IDs and TDs, again suggesting that the IDs decode hand actions through the typical AON. Furthermore, additional ANOVAs analyses performed to explore possible differences between the decoding of hand and foot action in the two groups in brain areas showing significant hand decoding by at least one of the two groups (Figure 3B; see details in supplemental results section) revealed only a small cluster in the left LOTC showing an interaction pattern compatible with a sensorimotor developmental effect (selectively weaker hand action decoding in IDs; at a lenient threshold of p = 0.01). The rest of the AON did not show such interaction.

**Figure 3.**
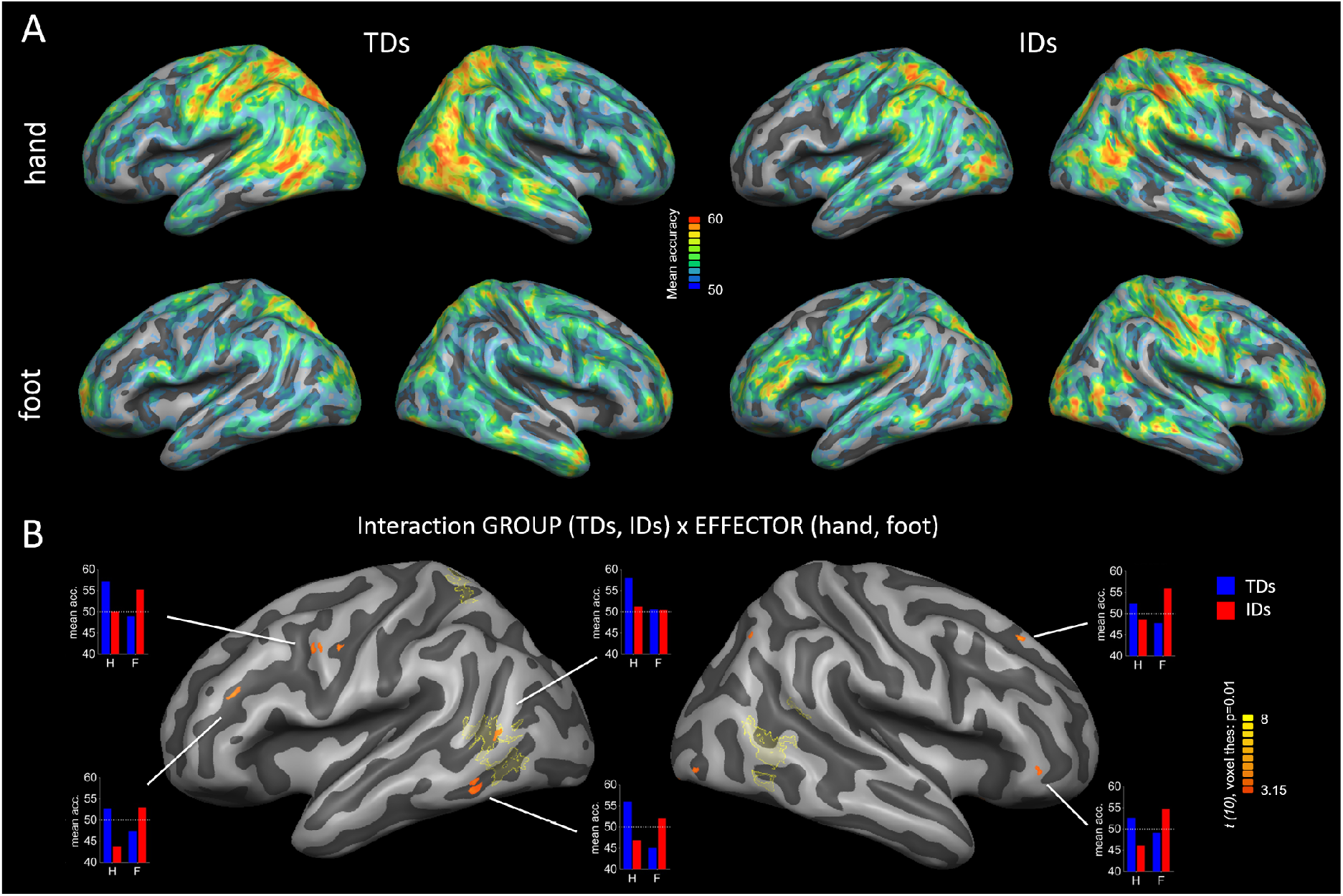
Across object decoding searchlight analyses. (A) Mean accuracy maps of hand an action decoding for the TDs and IDs show similar networks involved in hand action decoding i populations. (B) GROUP x EFFECTOR interaction map corrected using a group label bootstr procedure (see Methods for details). Yellow outlined areas indicate significant clusters of hand decoding of the TDs (corresponding to the TD mean accuracy maps in the upper left of pan corrected for multiple comparisons (voxel threshold p = 0.001, cluster threshold p = 0.05 interaction analysis shows few small clusters, only one of which (in the posterior left LOTC) area where decoding is significantly different from chance in the TDs. The mean decoding accu

#### Similarity of hand action decoding patterns

Since we found evidence for hand action decoding in the IDs and failed to find any substantial differences between the IDs and TDs, we performed additional ad hoc analyses to explore in more detail whether the networks subserving hand action representation are significantly similar in the IDs and TDs. To this end, we correlated the accuracy maps of the hand action decoding within the TDs and, then, between the TDs and the IDs. We reasoned that if the correlation between the two groups is (1) significantly above chance and (2) as strong as the within-TD correlation, then this would demonstrate that the action decoding patterns are similar in IDs and TDs. In a first step, we computed the mean within-TD correlation and compared it to the null distribution of correlations of random permutation maps of the TDs. The within-TD correlation was significant (above the 95% confidence interval of the within-TD random permutation correlations, p = 0.022), which suggests that the hand action decoding maps of the TDs contain significant information and are similar to each other (Figure 4A). Crucially, the between-group correlation was also significant (p = 0.006) and as strong (or stronger) as the within-TD correlation (Figure 4A).

**Figure 4.**
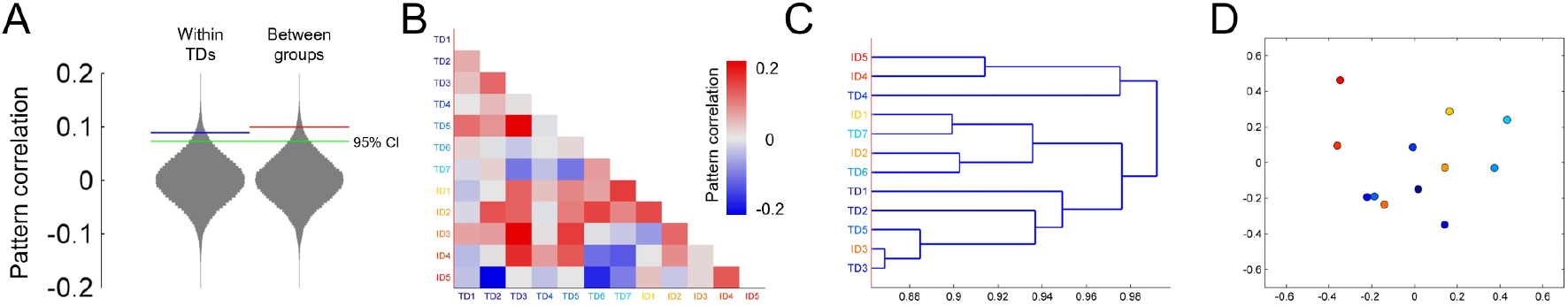
Pattern similarity of the across-object hand action decoding. (A) Mean within-TD correlation (blue line) and mean between group correlation (red line) superimposed on the distribution of within-TD random permutation correlations and its the 95% confidence interval (green line) show both within and between-group correlations are significant. (B) Correlation matrix of all participants’ decoding maps. (C, D) Hierarchical cluster analysis of the dissimilarities between the subjects’ maps (1 – r) visualized as dendrogram (C), and multidimensional scaling (D), both show the IDs maps fall within the variability of the TDs. Blue and red dots indicate TDs and IDs, respectively.

The result from these additional analyses thus provides positive evidence that the hand action decoding patterns of the IDs show significant similarity to the action decoding pattern of the TDs. To visualize the similarities of the individual action decoding patterns of IDs and TDs we correlated all subjects with each other (Figure 4B) and carried out a hierarchical cluster analysis. A hierarchical cluster analysis as well as a multidimensional scaling demonstrated that IDs and TDs did not form segregated clusters but were intermingled with each other (Figure 4C and 4D), again showing that the IDs action decoding maps are within the typical distribution.

### Across effector decoding

These findings invite the speculation that the LOTC and IPL could house action representations that are largely independent of the effector that is used to carry out the action (i.e., independent of whether an action is carried out with the hand or the foot). If so, these representations should generalize not only across the objects involved in the actions but also across the effector used for the actions. To test this possibility, we trained a classifier to discriminate between opening and closing an object with one effector (e.g., the foot) and tested the classifier to discriminate the two actions carried out with the other effector (e.g., the hand). We identified action representations that generalize across effectors as well as representations that generalize across both effectors and objects in the LOTC and IPL of both TDs and IDs (Figures S1 and S2). Moreover, exploratory group comparisons suggested that the patterns of these representations were more likely similar than different in TDs and IDs suggesting that also effector-invariant action representations are not substantially modulated by sensorimotor experience (Figures S1-S3; Table S3). Therefore, parts of the AON, including LOTC and IPL, may represent abstract action representations, beyond the used effector.

### Univariate analyses

Finally, we tested whether the IDs show deviant neural activations during the observation of hand actions. We thereby targeted representations that are not necessarily action-specific and/or feature-general, e.g. representations at processing stages that may precede the activation of high-level action representations. We found significant univariate activation in the IDs in the AON (Figure S4). In addition, traditional and Bayesian mixed ANOVAs and two-sample t tests performed to explore possible differences between the representation of hand and foot actions in the two groups suggested that the representation of hand and foot action are more likely similar than different between the groups (see supplemental results and supplemental tables S4 and S5). Finally, we found no evidence for hand action selective increase of activation in any other area of the brain (even at a liberal threshold) that could suggest the use of an alternative action processing mechanism. These results suggest a typical distribution of brain activation during the observation of hand actions in the IDs.

## Discussion

We used fMRI to study the processing of observed hand and foot actions in individuals born without upper limbs and, thereby, determine whether the typical large-scale organization of the AON is genetically determined or emerges through repeated effector-specific functional coupling within the bidirectionally connected functional circuit translating observed actions into the corresponding motor codes, and vice versa (the AON). Using a combination of univariate analyses, multivoxel pattern decoding, and pattern similarity analyses, we found the typical spatial arrangement and functional properties of brain areas specialized in the processing of observed actions for hand and foot actions both in participants born without upper limbs and in typically developed individuals. We additionally found no evidence for large group difference in the AON. This clear spatial and functional preservation for hand actions in the IDs, despite an absence of hand movement execution, demonstrates that effector-specific visuomotor coupling is not necessary for the areas belonging to the action observation network to specialize for observed human actions. Together with previous findings that specialization for hands and tools is also independent from sensorimotor experience (Striem-Amit et al. 2017), but not for letters (Baker et al. 2007; Dehaene et al. 2010; Saygin et al. 2016), our finding thus adds new evidence in support of a conception of brain organization that differentially attributes a primary role to genetic factors in the emergence of brain specialization for stimuli of evolutionary relevance (Caramazza and Shelton 1998), such as body parts, tools and human actions, and to post-natal experience for stimuli with less evolutionary relevance or that were acquired later during human evolution (such as letters and numbers).

This theoretical perspective does not imply that experience does not influence the content or fine-grained organization of the regions coding for evolutionarily relevant categories. Evidence that extensive training and experience can change the fine-grained response profile of the AON is compelling (Turella et al. 2013). Dancers, for instance, activate some parts of the AON more when they observe dance movements that they are used to perform than those they are less used to perform (Calvo-Merino et al. 2006). In the same vein, we also found a small cluster in the posterior dorsal left LOTC that showed selectively weaker hand action decoding in the IDs than in the TDs. Thus, the claim here is not that there are no subtle differences between groups or between the processing of hand and feet actions in the IDs. Our results suggest, however, that experience-dependent structural plasticity is highly constrained by genetically determined organization principles.

As such our findings corroborate those reported by two previous studies reporting on the brain correlates of action observation in individuals born without upper limbs. The first study reported on the brain correlates of observing actions (e.g., writing or crushing) that were possible or impossible to execute for a participant, DD, born without forearms and lower limbs (Aziz-Zadeh et al. 2012). The second study scanned two dysplasic subjects, born without arms or hands, while they watched hand actions (Gazzola et al. 2007). In both studies, the IDs’ AON was activated when they watched actions that they could not perform. Nevertheless, these observations were somewhat limited because the sole use of univariate analyses did not allow probing the representational content of the AON nodes in the IDs.

Beyond contributing to theories of the emergence of brain specialization, our findings are also relevant for theories about the brain substrate of action perception and interpretation. First, they extend previous studies by showing that the LOTC and IPL contain information about the content of an observed action that generalizes not only across viewpoint, kinematics, and the object involved in the action (Oosterhof et al. 2012; Wurm and Lingnau 2015; Wurm et al. 2016), but also across effectors regardless of sensorimotor experience. Second, our findings are also relevant for the question of the role of motor simulation in action recognition. According to motor simulation theories of action recognition, the recognition of others’ actions cannot be achieved by visual analysis of the movements alone but requires unconscious covert imitation – motor simulation – of the observed movements (Blakemore and Decety 2001; Rizzolatti and Sinigaglia 2010). At odds with this hypothesis, previous behavioral studies showed that the IDs perceive and comprehend hand actions, which they cannot covertly imitate, as accurately, as fast, and with the same biases as typically developed participants (Vannuscorps et al. 2013; Vannuscorps and Caramazza 2016, 2017). However, the possibility remained that the IDs use alternative, compensatory strategies or brain mechanisms to reach efficiency. The present findings suggest that this is not the case and, together with the previous behavioral findings, constitute clear evidence that efficient action recognition can be supported by the visual-cognitive brain structures unaided by the motor system.

In conclusion, the clear preservation of functional organization of the AON in people born without hands observing hand actions suggests that sensorimotor ontogenetic experience is not required for this specialization to emerge. Instead, it points to an evolutionarily driven functional selectivity, which can develop based on inherited connectivity constraints.

## Acknowledgements

We thank Himanshu Bhat and Thomas Benner of Siemens Healthcare for the SMS-EPI sequence, and Steven Cauley of Massachusetts General Hospital for modifications that enabled implementation of our protocols in a routine session. The authors declare no competing financial interests.

